# Genotype imputation in F2 crosses of inbred lines

**DOI:** 10.1101/2023.12.12.571258

**Authors:** Saul Pierotti, Bettina Welz, Mireia Osuna Lopez, Tomas Fitzgerald, Joachim Wittbrodt, Ewan Birney

**Affiliations:** European Molecular Biology Laboratory, European Bioinformatics Institute (EMBL-EBI); Cambridge, CB10 1SD, UK; Centre for Organismal Studies (COS), Heidelberg University; Heidelberg, 69120, Germany; European Molecular Biology Laboratory (EMBL), Genomics Core Facility; Heidelberg, 69117, Germany

## Abstract

**Motivation:** Crosses among inbred lines are a fundamental tool for the discovery of genetic loci associated with phenotypes of interest. In organisms for which large reference panels or SNP chips are not available, imputation from low-pass whole-genome sequencing is an effective method for obtaining genotype data from a large number of individuals. To date, a structured analysis of the conditions required for optimal genotype imputation has not been performed.

**Results:** We report a systematic exploration of the effect of several design variables on imputation performance in F2 crosses of inbred medaka lines using the imputation software STITCH. We determined that, depending on the number of samples, imputation performance reaches a plateau when increasing the per-sample sequencing coverage. We also systematically explored the trade-offs between cost, imputation accuracy, and sample numbers. We developed a computational pipeline to streamline the process, enabling other researchers to perform a similar cost-benefit analysis on their population of interest.

**Availability and implementation:** The source code for the pipeline is available at https://github.com/birneylab/stitchimpute. While our pipeline has been developed and tested for an F2 population, the software can also be used to analyse populations with a different structure.

## Introduction

Pure inbred lines and structured crosses have been the bedrock of early genetics (Mendel, 1866) and they have been used in the earliest organisms explored in the field (Morgan, 1910; Aida, 1921). Over time, phenotype-based markers have been replaced by anonymous DNA variation such as Single Nucleotide Polymorphisms (SNPs) (Lander, 1996). With the advent of cheap high throughput sequencing, one can use low pass sequencing of subsequent generations (e.g., F2 generation) to accurately determine the genotype of an individual. This process is a type of imputation (i.e. the statistical technique to infer missing data), and statistical models have been specifically developed for the linked inheritance patterns of DNA sequence variation. Imputation approaches have been used for plant breeding (Bhattarai *et al*., 2020), farm animal agricultural studies (Li *et al*., 2022) and model organism studies (Yao *et al*., 2021).

Most imputation methods in current use rely on a Hidden Markov Model (HMM) where the sample genotypes represent the emission space, and the reference panel haplotypes define a two-dimensional grid of hidden states where each row is a reference haplotype, and each column is a marker. This is known as the Li and Stephens model after their foundational work on statistical modelling of linkage disequilibrium and recombination rates (Li and Stephens, 2003). In populations where the founding haplotypes are known, for example F2 crosses from inbred mice, implementations of the Li and Stephens model such as R/qtl2 (Broman *et al*., 2019) can be used as an effective solution to the imputation problem. However, often the founding haplotypes are not available or too costly to obtain.

The success of genotype imputation in humans has been made possible by the construction of large reference panels of high-quality genotypes. Milestones in this direction are the HapMap Project (Altshuler *et al*., 2010), the 1000 Genomes Project (The 1000 Genomes Project Consortium, 2015), and the Haplotype Reference Consortium (McCarthy *et al*., 2016). Some of the imputation tools in widespread use that take advantage of reference panels are GLIMPSE (Rubinacci *et al*., 2021), QUILT (Davies *et al*., 2021), ShapeIT (Delaneau *et al*., 2019), Beagle (Browning *et al*., 2018), miniMAC (Fuchsberger *et al*., 2015), and Impute2 (Howie *et al*., 2009).

The reliance on reference panels or high-quality founder haplotypes has hindered the application of genotype imputation in less-studied organisms that lack extensive reference data. STITCH (Davies *et al*., 2016) overcomes this limitation by recognising that in populations founded by a limited number of individuals, the unknown ancestral haplotypes are sequenced at relatively high depth even though each single sample is sequenced at low depth. STITCH can impute genotypes from low-depth sequencing data by alternating the Li and Stephens HMM with an expectation maximisation (EM) step where randomly initialised ancestral haplotypes are tuned to maximise the expectation under the model of the observed sample genotypes. This alternating procedure is repeated a parameterised number of times, leading to the reconstruction of the founder haplotypes and the imputation of the sample genotypes. Several hyperparameters need to be optimised for the reconstruction to be optimal, key among them the number of ancestral haplotypes to be reconstructed (K parameter). This optimization can be performed either by relying on quality metrics that are internal to the model (info score) or on external validation (correlation of imputed and known genotypes). Although STITCH has been used successfully in a number of diverse populations (Nicod *et al*., 2016; Zan *et al*., 2019; Scott *et al*., 2021; Zha *et al*., 2023; Blain *et al*., 2024; Li *et al*., 2024), often the parameterisation and the assessment of the required level of coverage are presented as a single best choice and with little exploration of alternative parameters and experimental designs. Previous work explored some of the parameters that influence STITCH accuracy in an Angora rabbit population (Wang *et al*., 2022), but this work did not explore the complex interplay between sequencing depth, population size, and cross structure.

We are studying phenotypes in the medaka fish (*Oryzias latipes*), which is a small teleost fish native to Japan, Korea and eastern China (Wittbrodt *et al*., 2002). It has long been used as a model organism for genetic research thanks to its comparatively small genome (700.4 Mb) (Kasahara *et al*., 2007) and to its economic husbandry and tolerance to inbreeding. Taking advantage of these characteristics, an extensive medaka inbred panel has been established from a wild population (Fitzgerald *et al*., 2022). Inspired by some of the plant breeding strategies, we have been using 8 parental lines, in 10 different cross combinations, for phenotype mapping. A recent example, which we will use in this manuscript, generated a total of 2219 samples (of which 2177 were used in the final analysis).

A practical problem that we encountered was the optimal imputation of the DNA sequence of each individual. Given that our medaka lines exhibit some residual genetic variability, we reasoned that STITCH would be the ideal tool for this task. We optimised STITCH hyperparameters for this type of multi-parent cross by assessing the imputation accuracy with a sample of high-coverage genomes and we explored the optimal parameters for future cost-effective cross designs. We show that a correctly parameterised STITCH pipeline can effectively impute sample genotypes and show that there is a trade-off between the number of individuals sequenced and the per-sample coverage needed.

## Results

We first developed a framework for the optimisation and assessment of imputation results from F2 crosses. We performed high-coverage sequencing on one F1 individual per cross (10 overall) and two F2 individuals (from one cross only). We reasoned that we could use F1 individuals as high-coverage “truth sets” for the imputation assessment to maximise the probability of observing all the haplotypes from a given cross in the smallest possible number of validation samples. We show that this choice does not affect model selection and does not inflate performance in Supplementary Figure S3. The F1 individuals are also useful to determine the set of truly polymorphic markers in the cross. Although we have sequenced individuals from the F0 founder lines (Fitzgerald *et al*., 2022) both previous work (Jaegle *et al*., 2023) and our own investigations suggested that there will be apparent deviations from the expected inheritance patterns (e.g. in cryptic segmental duplications and in genomic regions that are not fully inbred), and so we preferred to develop a system where no assumptions about the founders’ haplotypes are made. Before using them for imputation, we downsampled the F1 and high-coverage F2 samples to obtain approximately equivalent coverage to the low-depth sequencing data across all F2s (Figure 1). Thus, we had 12 samples imputed from low-coverage data for which we have also a high-quality, high-confidence genotype call. Using this truth set we explored STITCH parameters.

**Figure 1.**
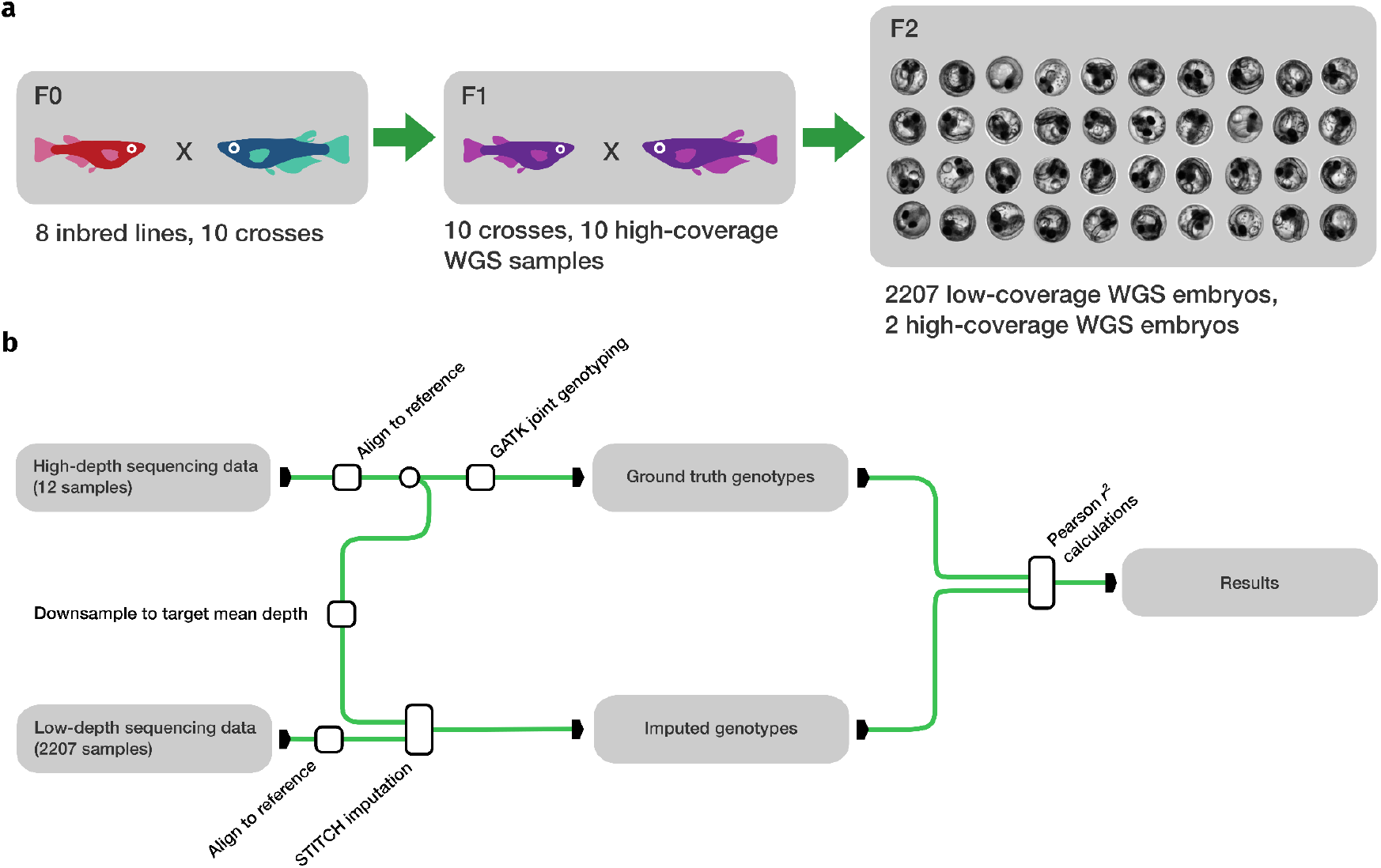
(a) Experimental design. 10 crosses of 8 inbred MIKK panel lines were performed. Among the resulting F1 progeny, 10 samples were sequenced at high coverage (see Methods). The F2 cohort was mostly sequenced at low coverage (2207 samples), but two F2 samples were sequenced at high coverage. WGS: Whole Genome Sequencing. (b) Schematic illustration of the analysis performed. High-depth samples were aligned to the medaka reference genome and jointly genotyped. Mapped reads were downsampled to the desired depth (see main text). Low-coverage samples were also aligned to the medaka reference genome. The low-coverage mapped reads and the mapped downsampled reads for the high-coverage samples were then jointly used for imputation with STITCH. The imputed genotypes for the high-coverage samples were compared to the ground truth genotypes that had been obtained with GATK joint calling for evaluating imputation performance (using the Pearson *r*^2^ as a metric).

In our hands, the two key parameters for a successful imputation were the number of ancestral haplotypes (K parameter) and the selection of valid SNPs to impute. To select the imputable SNPs, we started with the SNPs that are polymorphic in the 10 F1 individuals (one F1 per cross) and the two high-coverage F2s. We then refined this set by running the imputation including the downsampled high-coverage datasets, computing the correlation of the imputed SNPs to the high-coverage calls, and rejecting SNPs with a low correlation (Figure 2a). We repeated this process for 5 iterations, progressively increasing the stringency of the correlation threshold (see Methods). We show that this procedure substantially improves imputation performance and does not result in a biased dropout of low-frequency SNPs or SNPs present only in one founder line in Supplementary Figures S4, S5 and S6. For the selection of the best setting for K, we tested a range of values and selected the one that maximised the mean squared Pearson correlation across the 12 truth set samples (Figure 2b). As both K and the final SNP set parameters are linked, our procedure is to first determine a plausible value for K using the unfiltered SNP set, then doing the SNP filtering (using the value of K determined in the initial assessment), and finally re-assessing the choice of K on the filtered SNP set.

**Figure 2.**
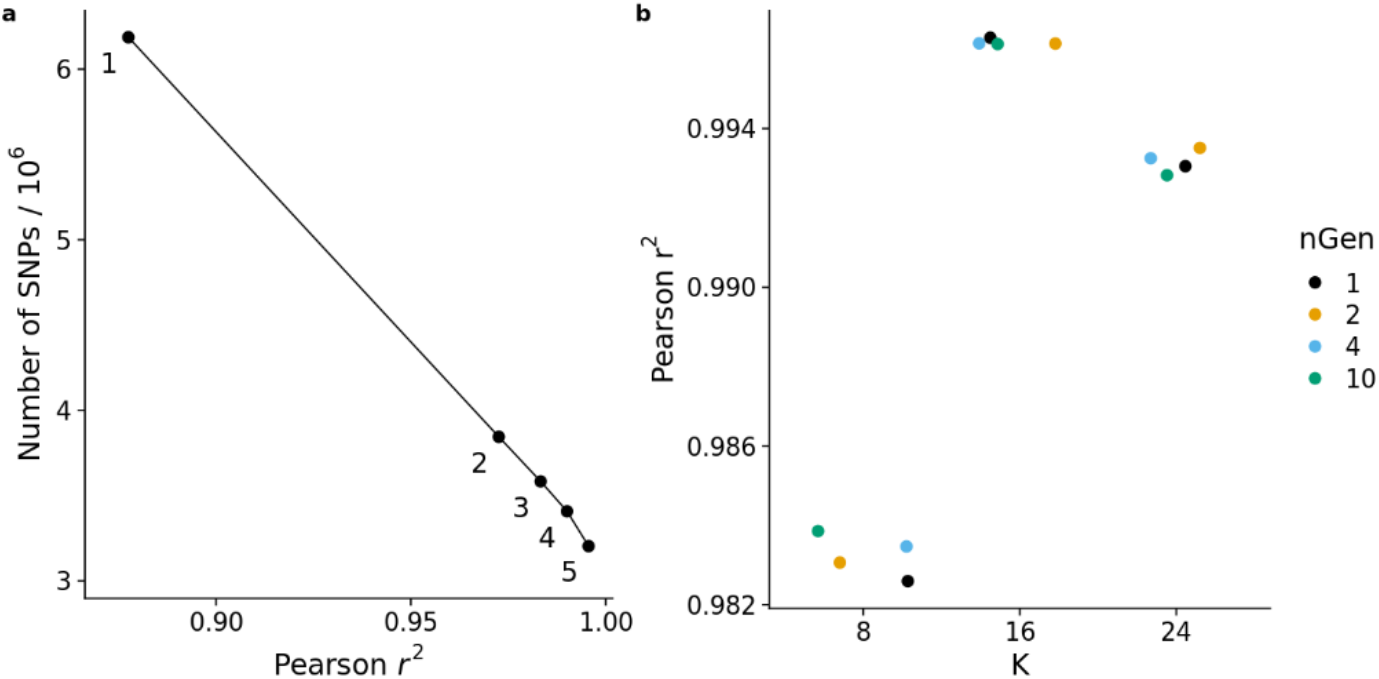
(a) Refinement of the imputed marker set showing the trade-off between the number of markers retained and imputation performance. The numeric labels refer to the refinement iteration. (b) Imputation performance for different hyperparameter combinations. K is the number of ancestral haplotypes and nGen is the number of generations since the founding of the imputed population. In our analysis, K=16 is optimal and nGen has only a modest effect on the results.

Overall, our F2-STITCH pipeline proved to be highly successful in imputing the genotypes of our medaka cohort. We observed a mean per-sample squared Pearson correlation coefficient (*r*^2^) of 0.996 between the imputed and ground truth genotypes. When binning the SNPs according to their Minor Allele Frequency (MAF), the squared Pearson correlation was as expected higher for bins containing more common SNPs (most likely due to being polymorphic in more than one cross) and decaying for rarer SNPs. Nonetheless, even the bin containing the least common SNPs (MAF 0 to 0.05) still showed an *r*^2^ of 0.977.

### Parameters that influence accurate imputation

Using the pipeline, we then explored downsampling our overall F2 dataset in a variety of dimensions: in the sequencing depth, in the number of individuals overall, in the number of crosses, and in the number of individuals in a single cross. This explores much of the design space for practical cross strategies.

As expected, we observed less accurate imputation for lower sequencing depth and lower sample numbers (Figure 3a, 3b), though there is a non-linear response in both cases. For example, we observed a large difference in imputation performance between a coverage of 0.5x and 0.25x, but a comparatively smaller difference when the coverage was changed from 1x to 0.5x. We observed a similar trend when reducing the total number of samples while retaining the original sequencing depth (Figure 3b). In this case, the performance started to drop significantly for cohorts that were smaller than 1000 samples.

**Figure 3.**
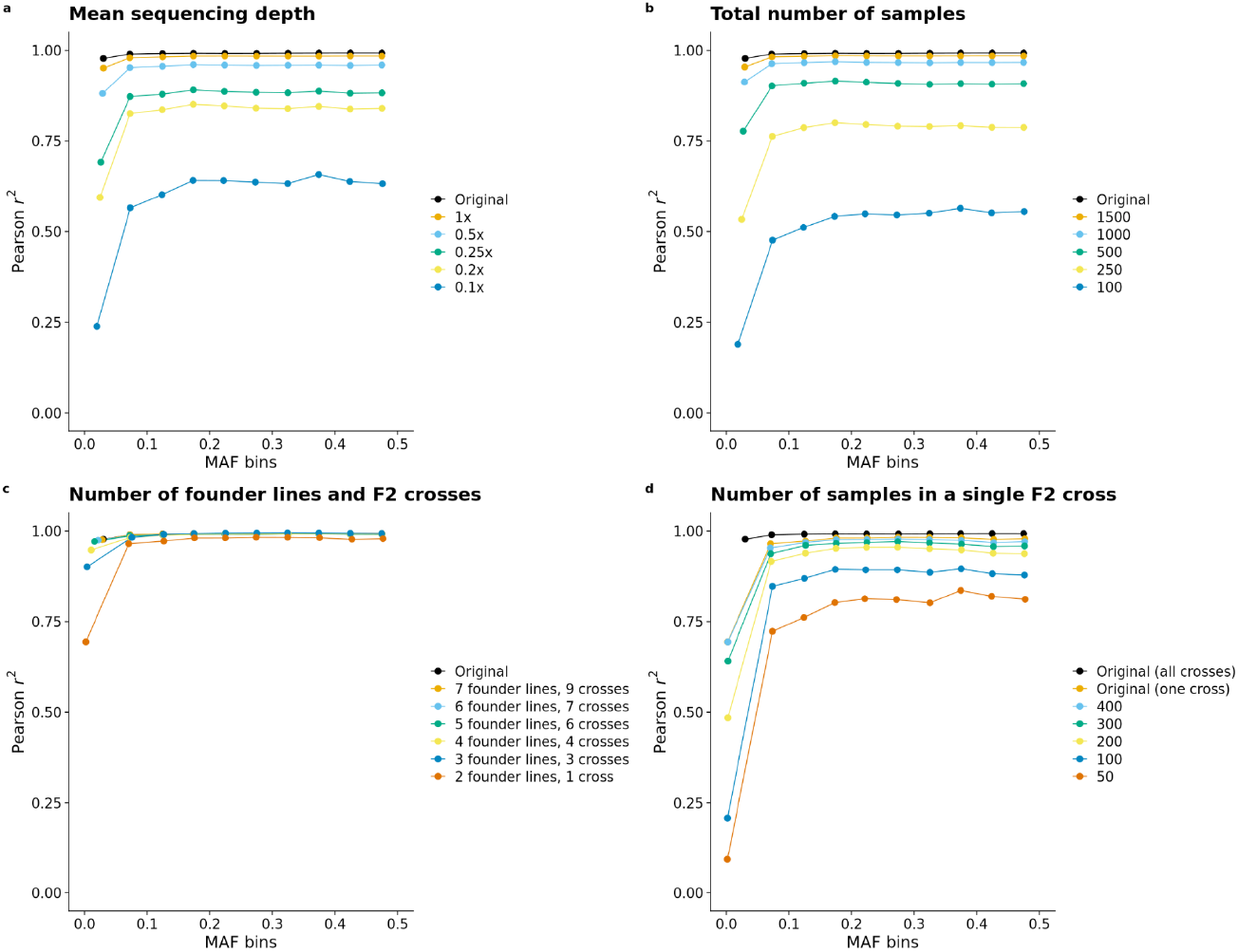
Imputation performance in different scenarios. All plots show the squared Pearson correlation for imputation compared to the truth set for different Minor Allele Frequency (MAF) bins. (a) Varying sequencing depth. (b) Varying the total number of samples jointly imputed. (c) Restricting the imputation to samples derived from a limited number of founder lines and F2 crosses. (d) Imputing a single F2 cross and varying the number of samples within that cross.

On the contrary, reducing the sample numbers by reducing the number of F2 crosses involved (and hence the number of distinct haplotypes to be reconstructed; we changed K proportionally, see Methods) did not significantly affect the quality of the imputation (Figure 3c). When the dataset was reduced to a single F2 cross (and so 2 F0 founder lines) for a total of 474 samples, the *r*^2^ value was observed to be above 0.964 for all the MAF bins except the one with the least common SNPs (which had a much lower *r*^2^ of 0.694). This compares to an *r*^2^ range of 0.777 to 0.915 when using 500 samples randomly chosen from the whole dataset (contrast Figure 3b, “500 random samples” to Figure 3c, “2 founder lines, 1 cross”, with 474 samples).

Finally, when we explored reducing the number of samples within a single F2 cross (Figure 3d), we observed that when the sample number fell below 200, the *r*^2^ sharply decreased. However, it should be noted that in a population similar to the one presented in this study, different F2 crosses share some common founder lines. As such, STITCH would be able to more accurately impute the genotypes compared to this simplified single-cross scenario by borrowing sequencing coverage of the founders from the other F2 crosses that have common parents.

### Cost-benefit analysis

To determine what would be the ideal sequencing depth to aim for when designing similar experiments, we performed a combined downsampling of mean sequencing depth and sample numbers. We observed that for the original cohort size (2177 samples) a maximum sequencing depth of 0.5x was sufficient for obtaining an imputation result that is only marginally worse than the original (Pearson *r*^2^ 0.981 versus 0.996) while almost halving the financial burden (53.7% of the original).

For lower sample numbers, the required minimum sequencing depth proportionally increased (Figure 4). For 1500 and 1000 samples, a depth of 1x was needed to avoid strong degradation of the imputation quality. For sample sizes lower than 500 any sequencing depth downsampling led to much worse imputation results. Also, we noticed that the Pearson *r*^2^ reached a plateau at different levels for different cohort sizes, with larger cohorts reaching the plateau earlier and ending at higher Pearson *r*^2^ values, though the difference between the best achievable correlations is marginal at high sample numbers.

**Figure 4.**
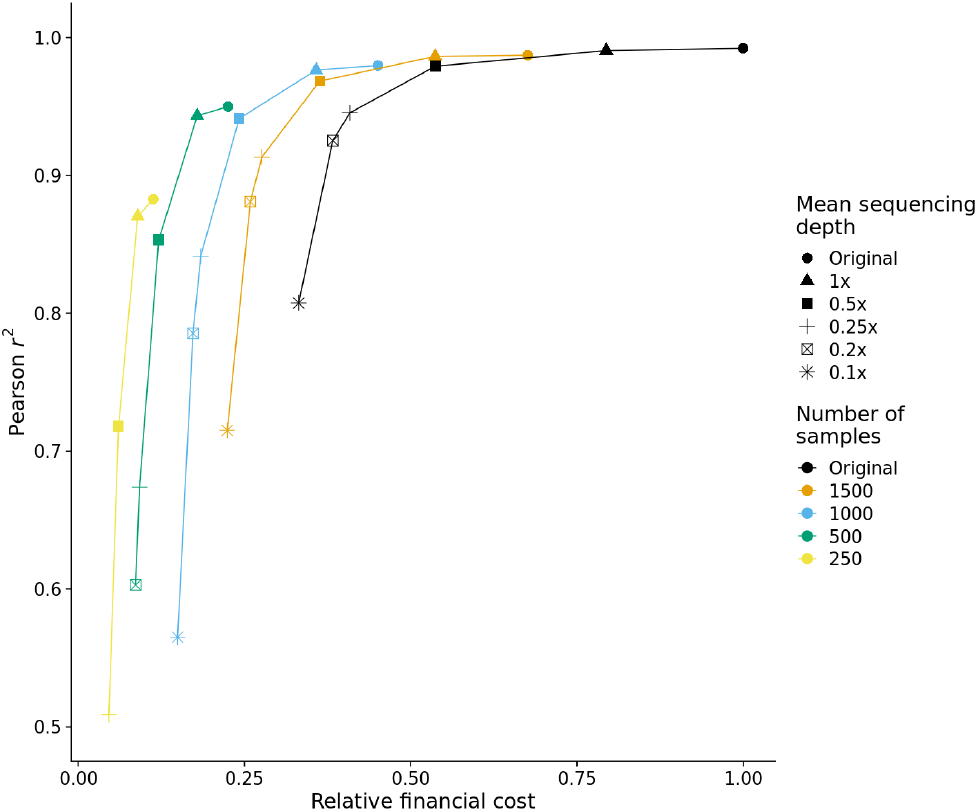
Relative cost of library preparation and sequencing versus imputation accuracy for different combinations of mean sequencing depth and number of samples. Estimated costs are relative to the cost sustained for obtaining the original dataset (see Methods). Depending on the number of samples, the imputation accuracy reaches a plateau at a certain level of mean sequencing depth, where increasing the depth further does not bring appreciable increases in imputation accuracy.

## Discussion

In this work, we have systematically explored the imputation accuracy from low-coverage sequencing for the commonplace F2 cross designs used in agriculture, model organisms and plant breeding. We also developed an economic model for the sequencing cost, on the assumption that the sample numbers would be fixed by other constraints (e.g. husbandry capacity). There is a relationship between the optimal sequencing depth and the number of samples for accurate imputation, as expected by the coverage of the underlying ancestral haplotypes. In practice, for sample sizes of 1000 aiming for a coverage of 1x at least for this cross structure provided nearly equivalent accuracy compared to our full set. In contrast for 2,000 samples a similar level of accuracy would be achieved with 0.5x. An alternative way of viewing this is that for sample sizes of 2000, there is little to be gained by raising the sequencing depth from 0.5x to 1x.

However, it is important to note that the optimal coverage and sample size cannot be determined *a priori* for an arbitrary population. Our results shall serve only as a guideline for similar experimental designs, as the genetic diversity and homozygosity level of the founders will have an impact on the optimal experimental design. In general, more diverse populations will require higher levels of coverage and/or larger sample sizes (see Supplementary Notes). Nonetheless, our pipeline can be used for exploring context-specific experimental and software parameters in such scenarios.

An important feature in both the practical F2 imputation and our assessment was the sequencing of F1 individuals at high coverage. In our experience, this was critical to select an initial set of SNPs which were polymorphic in the crosses and to assess imputation accuracy for different parameter settings. In practice, this means it is important to track and store F1 individuals during the breeding. As not all inbred lines are fully inbred, it is also advantageous to set up F2 crosses between single F0 individuals and track the relationship between F2 individuals and their specific F0 parents wherever possible. A benefit of STITCH is that it does not require a reference panel and will impute variants which are present in the F2 regardless of whether they were fixed in the F0 or not. While the info score produced by STITCH could be used for performance assessment in place of an external validation set to reduce cost, we suggest using an external validation when possible (see Supplementary Notes).

Another point to note is that we used a different DNA extraction and sequencing library preparation strategy for the high sequencing coverage and low sequencing coverage samples (ligation-based and PCR-free in the first case, tagmentation-based in the second). Tagmentation-based libraries are cost-effective for large numbers of samples, while ligation-based PCR-free workflows are free from transposase binding site bias and PCR artefacts, and so are more suitable for obtaining a high-quality validation set. While this differential processing could introduce systematic differences in sensitivity and precision in detecting variants (Ribarska *et al*., 2022), we think it does not substantially influence the parameter selection and coverage/sample size assessment performed in this study because: 1) a predetermined set of variants is used for imputation, and so false positives in the low-coverage samples have no impact on the imputation process; 2) the high-coverage samples are only used to validate the imputation process, and are not directly compared to the low-coverage samples; 3) the relative imputation accuracy between two imputation runs depends on the respective ancestral haplotype reconstructions, which are the same for both sets of samples. However, it must be noted that the *r*^2^ values reported in this work refer to the high-coverage samples only and may not be fully representative of the imputation performance in samples prepared with a different methodology.

A consequence of imputing only the highly confident polymorphic SNPs is that many other variants will not be imputed, including copy number variants, structural variants, insertions, deletions and hard-to-call SNPs. This does not invalidate the accurate imputation of the markers we have, but for downstream applications such as association studies, analysts should remember that the causal variant is not necessarily among the imputed set of markers. In our hands, a successful exploration technique is to group samples by the associated marker of interest and treat the grouped samples as a high-coverage meta-sample to explore other forms of genetic variation linked around a selected marker.

We have made this STITCH-based imputation scheme into a Nextflow (Tommaso *et al*., 2017) pipeline inspired by the nf-core template (Ewels *et al*., 2020) and available at https://github.com/birneylab/stitchimpute. This pipeline can take as inputs high-coverage and low-coverage samples, with a specification of any sequencing depth downsampling to be performed (e.g. for later assessment) and will automatically iterate the imputation and performance assessment for building a final SNP set. The pipeline also allows the exploration of the STITCH parameters K (number of haplotypes) and nGen (number of generations) in a systematic manner. It can also be used in a more basic way to run a single imputation with a pre-defined SNP and parameter set. We have successfully used this pipeline in a number of F2 crosses with different operators and hope it will be useful to a broader set of research groups. We are currently processing a number of phenotypes on this and other complex cross designs and are aiming for a comprehensive association study using these imputed genotypes.

Although we have explored parameters and downsampling for this particular cross structure in this specific species (medaka fish), different cross structures with different divergence levels of parents and different recombination rates will mean that some level of parameter adjustment will be needed for each system. Our recommendations are: (a) to ensure that there are high-coverage F1 individuals (b) to carefully consider the sequencing depth compared to the number of samples in the F2 and (c) to consider the number of parents and the number of founding haplotypes.

One aspect that we did not explore in detail in this work is the definition of the initial set of SNPs used in the imputation and filtering process. A possible extension that could be implemented in a future version of our pipeline is the use of a variant caller designed for (ultra) low-coverage sequencing data such as BaseVar (Liu *et al*., 2023) for defining this initial variant set without relying on the high-coverage samples.

## Materials and methods

### Fish maintenance

All fish stocks were maintained and bred (fish husbandry, permit number 35–9185.64/BH Wittbrodt) in constant recirculating systems at 26 °C on a 14 hrs light/10 hrs dark cycle. Fish husbandry and experiments were performed in accordance with local animal welfare standards (Tierschutzgesetz §11, Abs. 1, Nr. 1) and with European Union animal welfare guidelines (Bert *et al*., 2016). The fish facility is under the supervision of the local representative of the animal welfare agency.

### DNA extraction

The genomic DNA of the F1 individuals was extracted from whole brain samples. Brains were digested overnight at 60°C with proteinase K (1 mg/ml) in DNA extraction buffer (0.4 M Tris/HCl pH 8.0, 0.15 M NaCl, 0.1% SDS, 5 mM EDTA pH 8.0). Subsequently, the DNA was purified by phenol-chloroform isoamyl alcohol (Phenol:Chloroform:Isoamyl (25:24:1) Ph 8.0) extraction using phase lock gel (PLG) tubes. After centrifugation, the aqueous phase was mixed with Chloroform: Isoamyl Alcohol (24:1) in a new pre-spun PLG tube and centrifuged again. The genomic DNA was precipitated with Isopropanol and washed twice with 70% ethanol. The precipitated DNA was resuspended in TE buffer (10 mM Tris pH 8.0, 1 mM EDTA in RNAse-free water) and stored at 4 °C. For the genomic DNA extraction of the F2 individuals, hatched or manually dechorionated embryos were snap-frozen and stored in deep well plates at −80°C. The fish were lysed in 40 µl DNA extraction buffer (2M Tris pH 8.0, 5M NaCl, 20% SDS, 0.5M EDTA pH 8.0) overnight at 60°C. After a 1:2 dilution of the samples with nuclease-free water, the samples were used for library preparation.

### Sequencing library preparation

For the high sequencing coverage samples, a ligation-based library preparation was performed using the NEBNext® Ultra™ II DNA Library Prep Kit (New England Biolabs) with NEBNext® Multiplex Oligos for Illumina® (Unique Dual Index UMI Adaptors DNA Set 1) according to the manufacturer’s instructions without PCR. 750 ng DNA were used as input. The libraries were selected for a fragment size of 500 bp and quantified with the Qubit High Sensitivity kit. Further, the quality and the molarity of the library were assessed using the Bioanalyzer with the DNA HS Assay kit following the manufacturer’s protocol allowing equimolar pooling of libraries for sequencing. For the low sequencing coverage samples, a tagmentation-based library was prepared. The sequencing library preparation was done as described previously (Picelli *et al*., 2014). For the tagmentation reaction, 1,25 µl DNA was mixed with 1,25 µl of Dimethylformamide, 1,25 µl tagmentation buffer (40 mM Tris-HCl pH 7,5, 40 mM MgCl2) and 1,25 µl of an in-house generated and purified Tn5 (Hennig *et al*., 2018) and incubated at 55°C for 3 min. To inactivate the tagmentation reaction the samples were cooled down to 10°C and 1,25µl 0,2 % SDS was added followed by an incubation for 5 min at room temperature. The resulting fragments were amplified by PCR (72°C for 3min, 95°C for 30sec, 12 cycles of 98°C for 20 sec, 58°C for 15 sec and 72°C for 30 sec and a final extension time of 3 min at 72°C) using 6,75 µL of KAPA 2 X HiFi master mix, 0,75 µL of Dimethyl sulfoxide and 2,5 µL of dual indexed primers. PCR products were pooled and selected for an average size of 450 bp using two rounds of magnetic SPRI beads cleanup (0.7X and 0.6X respectively). Finally, the library was quantified with the Qubit High-Sensitivity kit and the fragment size was assessed using the high-sensitivity assay in the Bioanalyzer.

### Sample exclusions

Samples sequenced to an overall mean depth lower than 0.1x were excluded from the analysis. This reduced the sample set from 2219 to 2177 samples.

### F2 cross design

The population that we designed was founded by eight inbred medaka lines from the Medaka Inbred Kiyosu Karlsruhe panel (MIKK panel) (Fitzgerald *et al*., 2022). The lines used were 72-2, 55-2, 139-4, 15-1, 62-2, 68-1, 79-2, and 22-1. Those lines were crossed in a set of 10 F2 crosses (72-2 x 79-2: 153 samples, 72-2 x 15-1: 141 samples, 72-2 x 139-4: 153 samples, 139-4 x 72-2: 149 samples, 72-2 x 55-2: 481 samples, 72-2 x 62-2: 142 samples, 72-2 x 68-1: 164 samples, 68-1 x 79-2: 147 samples, 15-1 x 62-2: 158 samples, 55-2 x 139-4: 373 samples, 72-2 x 22-1: 148 samples). The cross 72-2×139-4 was performed reciprocally with males from the 72-2 line and females from the 139-4 line, and vice-versa. The other crosses were performed with males from the first line in the cross name, and females from the second line. One male and multiple females were used for founding each cross.

### Sequencing

A total of 2219 medaka samples were sequenced at the Genomics Core facility of the European Molecular Biology Laboratory (EMBL) in Heidelberg, Germany (https://www.embl.org/groups/genomics). We performed 150 bp paired-end Illumina (https://www.illumina.com/) short-read sequencing on a NextSeq2000 machine, with a variable number of samples per flow cell. In total, we sequenced our samples on nine P3 Illumina flow cells and two Illumina P2 flow cells. 12 samples were sequenced at high depth (33x to 61x) to be used as a ground truth for validating the quality of the imputation. Of these, 10 were F1 samples (one per cross, multiplexed on a P3 flow cell) while the remaining two were F2 samples (multiplexed on a P2 flow cell). The 2207 low-coverage samples were sequenced at lower depth (1.4x overall and 9.5x maximum per sample) on eight P3 flowcells (multiplexed with 271, 276, 276, 287, 288, 359, 90, and 90 samples respectively) and one P2 flow cell (270 samples). The mean per-sample sequencing depth distribution for all the samples is reported in Figure S1.

### Alignment and preprocessing

Both low-coverage and high-coverage (ground truth) samples were pre-processed using the nf-core/sarek (Hanssen *et al*., 2023) pipeline. Reads were mapped to the HdrR medaka reference genome (ENSEMBL ID: ASM223467v1) using bwa-mem2 (Vasimuddin *et al*., 2019) and deduplicated using GATK MarkDuplicates (Van der Auwera and O’Connor, 2020).

### Sequencing depth calculations

All the sequencing depths reported in this work have been calculated on the output of GATK MarkDuplicates using Mosdepth (Pedersen and Quinlan, 2017), as part of the nf-core/sarek pipeline. The coverage was obtained from the Mosdepth summary output by summing the total number of bases sequenced for each chromosome and dividing by the sum of chromosome lengths. The mitochondrial chromosome was excluded from this calculation.

### Sequencing depth downsampling

The original mean depth for each sample (*d*_*original*_) was calculated as described above. Given a desired downsampled mean depth (*d*_*target*_), the downsampling factor (*f*) to be used was calculated as *f* = *d*_*target*_ / *d*_*original*_.

Samtools view (Danecek *et al*., 2021) was then used to downsample the reads with the calculated downsampling factor using the --subsample flag.

Given the variable original mean depth of the low-coverage samples, not all the samples always had an original depth higher than the target depth. In such cases, no downsampling was performed. As a consequence of this, the depth values indicated represent an upper bound on the real mean depth across the cohort.

When assessing the imputation performance on the dataset with original sequencing depth, the ground truth samples were downsampled to a mean depth of 0.5x before being imputed, so as to avoid inflated performance figures driven by the relatively high coverage. When the depth was downsampled, the ground truth samples were also downsampled at the same target depth used for the rest of the cohort. Nonetheless, we observed that the specific coverage of the ground truth samples had a negligible effect on the imputation performance, while the overall coverage of the imputed cohort was much more important.

### Sample size downsampling

The number of samples was downsampled in such a way that the ground truth samples were always retained. Among the remaining samples, the fish to be included were chosen randomly until the sample set (complete of ground truth samples) reached the desired size. In the overall downsampling, samples were chosen randomly without considering F2 cross membership (Figure 3b).

### F2 crosses downsampling

In the downsampling performed in Figure 2c, all the samples belonging to certain F2 crosses were removed from the sample set (including ground truth samples belonging to such crosses). The F2 crosses and founder lines were iteratively downsampled in the following order: 1) start from the full sample set (8 founders, 10 crosses); 2) remove all the crosses involving line 22-1 (7 lines, 9 crosses); 3) remove all the crosses involving line 68-1 (6 lines, 7 crosses); 4) remove all the crosses involving line 79-2 (5 lines, 6 crosses); 5) remove all the crosses involving line 15-1 (4 lines, 4 crosses); 6) remove all the crosses involving line 62-2 (3 lines, 3 crosses); 7) remove all the crosses involving line 139-4 (2 lines, 1 cross).

### Single F2 cross downsampling

The single cross downsampling in Figure 3d starts from the subset remaining at the end of the F2 cross downsampling in Figure 3c. This sample set includes only the samples belonging to the cross 72-2 x 55-2. The samples in this cross were then randomly selected while retaining the ground truth samples, until the desired sample size was reached.

### Ground truth genotyping

The 12 ground truth samples were genotyped using the nf-core/sarek pipeline. The reads were aligned to the reference genome using bwa-mem2 and deduplicated using GATK MarkDuplicates. Genotypes were obtained with GATK via joint germline variant calling (Poplin *et al*., 2018).

### Imputation

Imputation was performed using STITCH (Davies *et al*., 2016). The following parameters were used: K=16, nGen=2, expRate=2, niterations=100, shuffleHaplotypeIterations=seq(4, 88, 4), refillIterations=c(6, 10, 14, 18), shuffle_bin_radius=1000. The K parameter was chosen so as to maximise the squared Pearson correlation with the ground truth on the full medaka genome. The other parameters were set by inspecting diagnostic plots produced by the software when imputing a small genomic region in an initial exploratory phase of the project. These other parameters were determined to have a far smaller impact on imputation accuracy than the K parameter.

When downsampling the number of crosses and founder lines, the K parameter was adjusted proportionally to reflect the fewer number of founder lines present in the population. Consistently with the choice of K=16 determined to work best in the original dataset that was composed of 8 founder lines, K was set to twice the number of founder lines in each downsampling iteration. When downsampling the number of samples in a single cross, K was set to 4 to reflect the presence of 2 founder lines.

A graphical representation of the ancestral haplotype usage along the genome for the optimal parameter set is reported in Figure S2.

### SNP set definition

The set of SNPs used in the imputation was defined iteratively. We started from a set containing all the biallelic SNPs called by GATK joint calling in the ground truth sample set with a minor allele count greater or equal to 1 (6.2 million SNPs). To obtain this we used bcftools view (Danecek *et al*., 2021) and applied in series the following filtering criteria: -i ‘CHROM!=“MT”’, -i ‘TYPE==“snp”’, -i ‘N_ALT >= 1’, -I ‘MAC >= 1’. We ran the imputation jointly on the low-coverage and downsampled ground truth samples (0.5x mean depth) and measured the SNP-wise squared Pearson correlation against the ground truth genotypes. We retained SNPs only above a certain threshold of squared correlation and then repeated the imputation on such SNP set. We iteratively refined the SNP set in this way for a total of 5 iterations, using filter values of 0.5, 0.5, 0.75, 0.9, and 0.9. The final SNP set consisted of 3.2 million SNPs. This filtering procedure follows the approach suggested in the original STITCH publication (Davies *et al*., 2016).

### Imputation quality calculations

The quality of the imputation was calculated in terms of squared Pearson correlation (*r*^2^) between the imputed genotypes and the genotypes obtained via joint germline GATK calling on the ground truth samples. GLIMPSE2 Concordance (Rubinacci *et al*., 2021) was used for this task. The performance was calculated jointly for SNPs in Minor Allele Frequency (MAF) bins of size 0.05. The plotted MAF of each bin represents the average observed MAF of the SNPs that it includes. The performance was also calculated jointly for all the SNPs of a certain ground truth sample. This per-sample performance was averaged across samples to obtain single performance figures per imputation run like the one used in Figure 3.

### Relative cost calculations

The relative cost estimates reported in Figure 3 represent the fraction of the original cost that would have approximately been spent for a certain downsampling scenario. These were calculated from the sequencing and library preparation cost at the Genomics Core facility of the European Molecular Biology Laboratory (EMBL) in Heidelberg, Germany. The following formula was used to compute the cost: ((*n ∗ d ∗ g*) / *f*) *∗ c*_*f*_ + *n ∗ c*_*p*_.

In the above formula, n is the number of samples, d is the mean sequencing depth, g is the haploid genome size, f is the mean output per sequencing flow cell, and *c*_*f*_ and *c*_*p*_ are the unitary costs for sequencing (per flow cell) and library preparation (per sample).

The mean output per flow cell was estimated from the mean observed output in our experiment for P3 Illumina flow cells (*f* = 273542732637 *bp* after deduplication). The medaka haploid genome size (*g* = 734040372 *bp*) was obtained by summing the chromosome lengths of the Hdr-R reference genome, mitochondrion excluded. The unitary costs had a ratio *c*_*f*_ / *c*_*p*_ = 683.89.

From this, the relative cost was computed by dividing the result by the cost estimated using the number of samples and the mean depth of the original sample set.

### Additional software used

Plots were produced using ggplot2 (Wickham, 2016) and cowplot (Wilke, 2020). JupyterLab was used for interactive explorations (Kluyver *et al*., 2016). The R programming language was used throughout the pipeline (R Core Team, 2023).

## Acknowledgements

We thank all the Birney and the Wittbrodt labs members for their support and constructive feedback on the work and the manuscript. We wish to acknowledge Amos Pierotti for providing the medaka fish illustrations used in Figure 1. We thank Vladimir Benes, Ferris Jung and all members of the EMBL Genomics Core Facility (GeneCore) for their support in the library preparation and the whole genome sequencing. This research was supported by the European Research Council Synergy Grant IndiGene [grant number 810172].

## Supplementary notes

### Ideal sequencing depth and sample size in different populations

Our results indicate that a minimum sequencing depth of 0.5x is required for 2177 samples from our complex F2 cross setup in medaka fish. In Davies *et al*., 2016 a similar sequencing depth versus sample size analysis is performed on 2073 CFW mice (Nicod *et al*., 2016) (outbred population established from 2 founders) and on 11670 Han Chinese women from the CONVERGE study (Cai *et al*., 2015). For the CFW mice, the first noticeable drop in imputation performance in the full sample set can be observed between 0.06x and 0.09x. For the CONVERGE study, the sample size does not seem to have a large impact on performance. For a sample size of 2000, a noticeable difference in correlation can be observed when moving from a sequencing depth of 1.4x to a sequencing depth of 1x.

This showcases how a more diverse population (CONVERGE study) requires higher sequencing depth for good imputation, while a less diverse population (CFW mice) can be reliably imputed with a lower sequencing depth. From our observations in terms of required sequencing depth, our medaka population is somewhat intermediate between these 2 examples.

### Use of the info score for filtering variants

We tested whether the info score provided by STITCH can be a suitable replacement for the external validation approach based on high-coverage sequencing that we used. We observe that the rank-based Spearman correlation between the info score and the external validation *r*^2^ is 0.58 for the full SNP set prior to filtering, and 0.51 for the refined SNP set after iterative filtering. These values were calculated excluding sites for which the validation *r*^2^ was missing (for example, sites that are not polymorphic in the ground truth samples). The total number of retained sites was 6.2 million for the unfiltered set and 3.1 million for the filtered set. The relatively low correlation between the info score and the external validation performance indicates that external validation is to be preferred whenever possible.

## Supplementary figures

**Figure S1.**
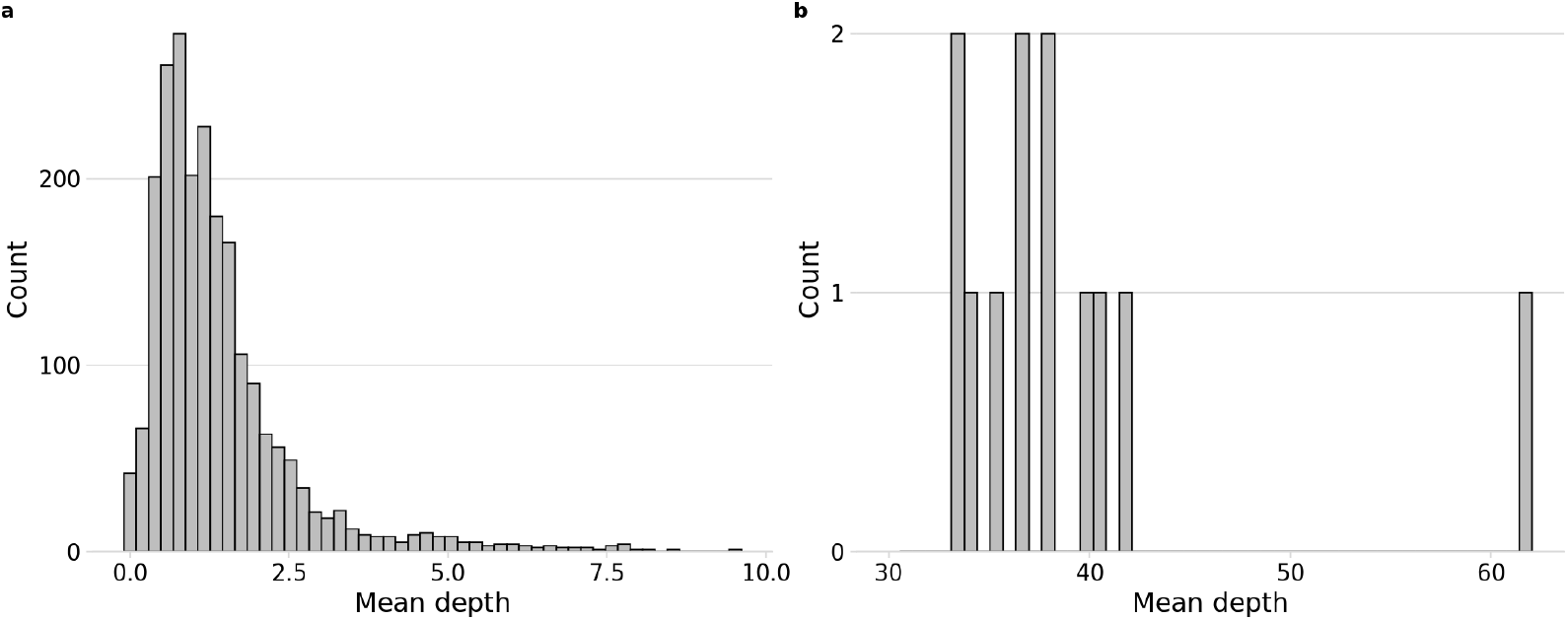
Distribution of mean sequencing depths in the original dataset used in this study. (a) Low-coverage samples. (b) Ground truth samples.

**Figure S2.**
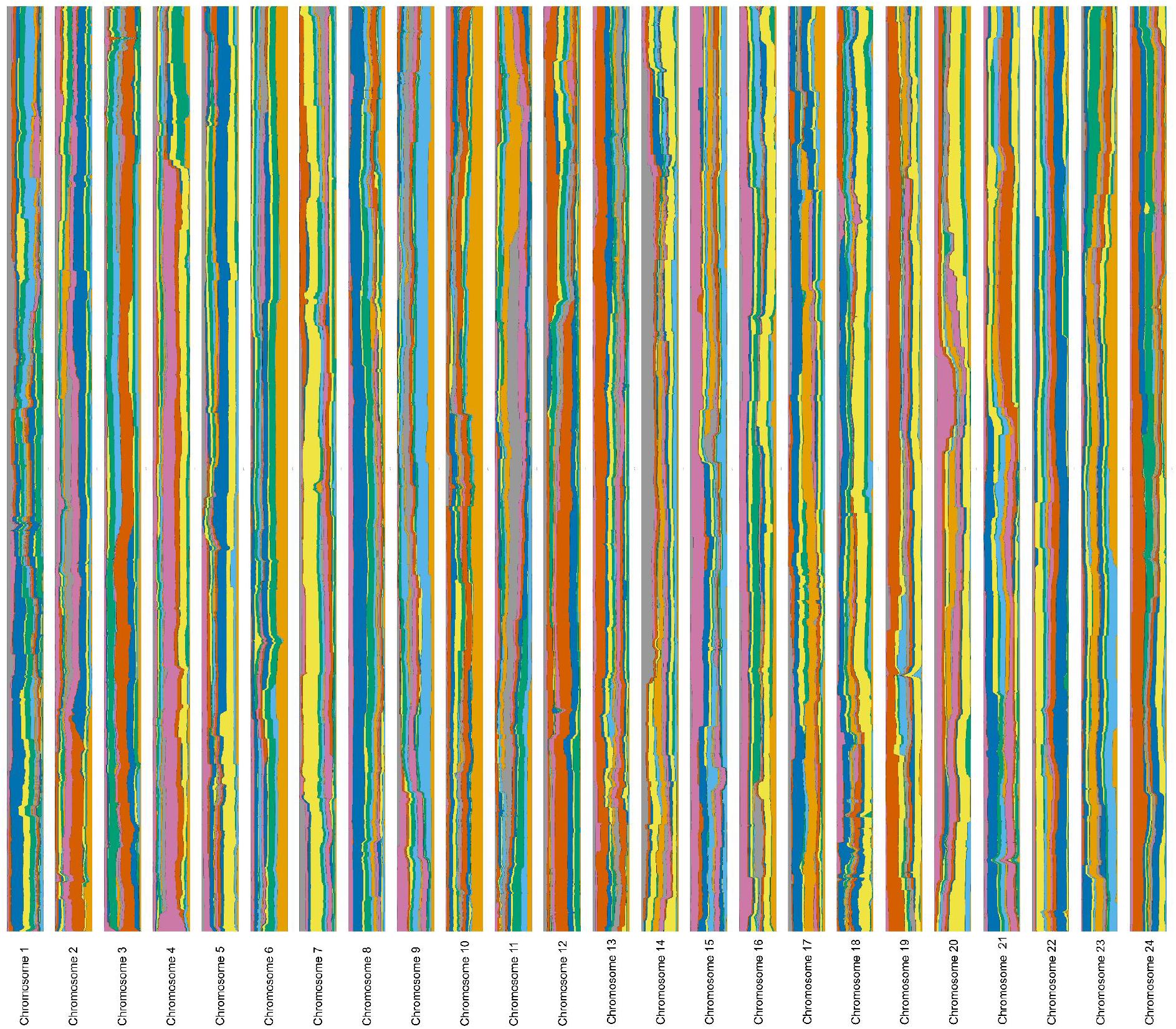
STITCH haplotype usage along the different chromosomes for the imputation run with the truth set samples downsampled to 0.5x mean depth and all the other samples left at the original depth. This visualisation has been produced by joining the per-chromosome plots automatically produced by STITCH. Long segments of contiguous colour reflect a lower number of ancestral haplotype switches, and hence a better imputation. Approximately equal usage of all the haplotypes also reflects more optimised heuristics.

**Figure S3.**
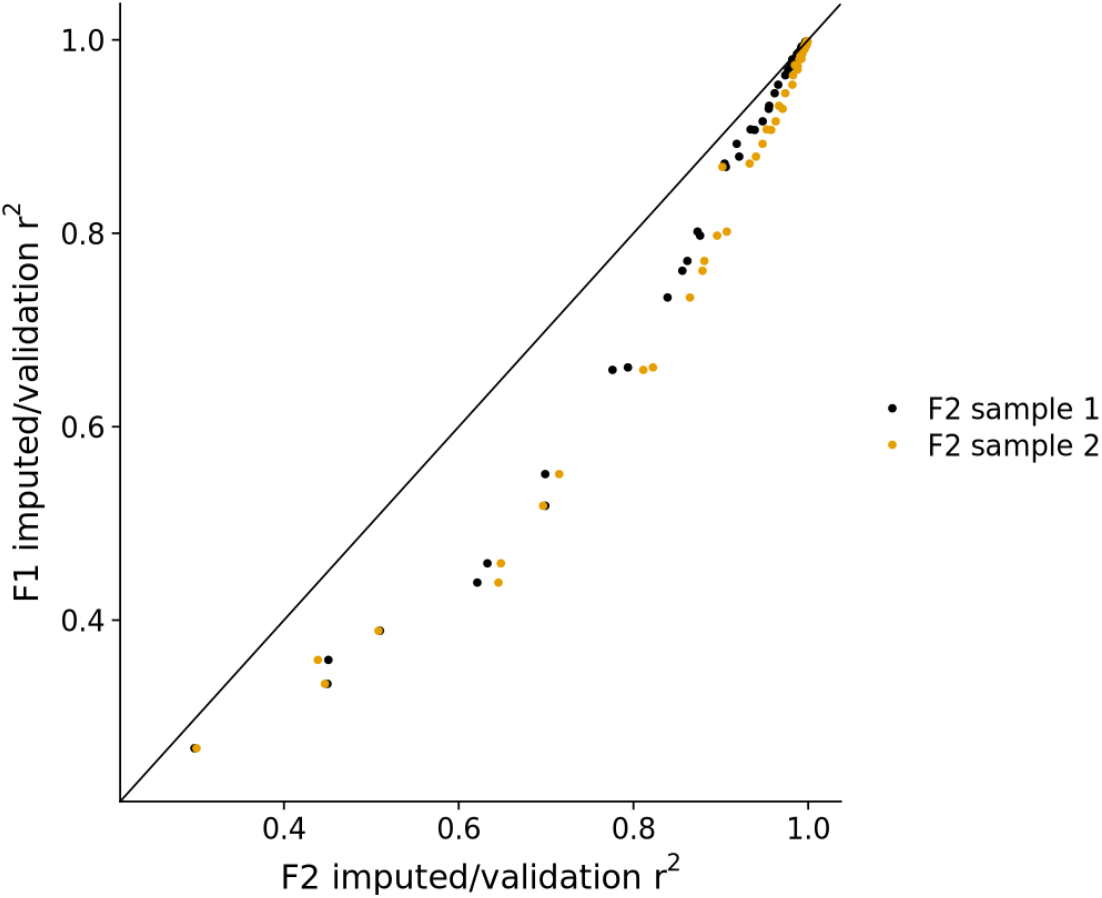
Comparison of the imputation performance evaluation on F1 samples and F2 samples from the same cross. For the F2 cross 72-2 x 55-2, we sequenced at high coverage one F1 fish and two F2 fish. This plot compares the sample-wise performance metrics for all the imputation runs performed in this work (each point is an imputation run) evaluated on the F1 sample (y-axis) and on the F2 samples (x-axis, different colours represent different F2 samples). F1 sample performance is an underestimation of the performance on F2 samples at intermediate correlation levels, while it tends to agree with the F2 performance towards the boundaries (0, 1). Importantly, the ranks of different performance runs are preserved (Spearman rank correlation = 0.99 for both F2 samples against the F1 sample), and so the F1 performance can be safely used for model selection in place of F2 performance.

**Figure S4.**
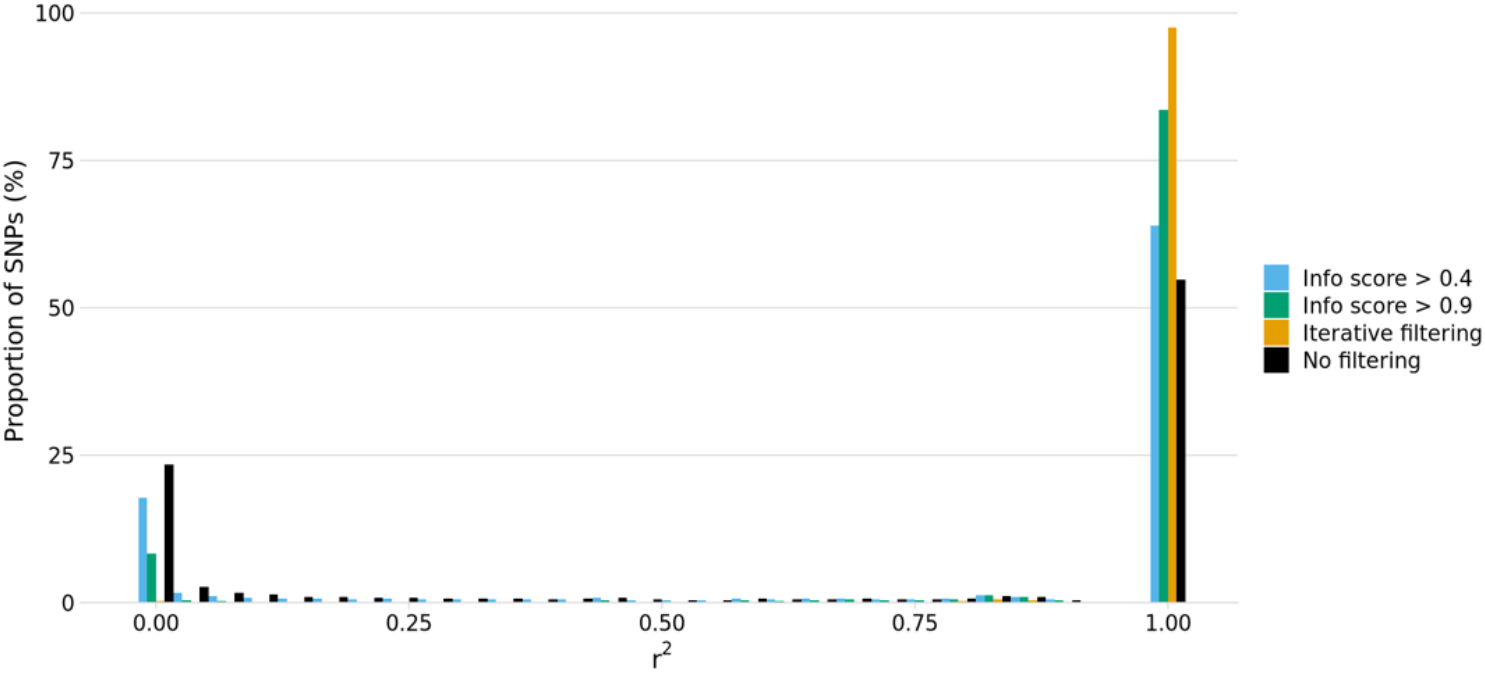
Distribution of SNP-wise imputation performance using different SNP filtering strategies. Iterative filtering is the approach that we used in this work, based on iterative imputation and filtering based on a comparison to the ground truth. The info score is an imputation quality metric internally produced by STITCH. Before filtering, many SNPs with very low imputed/validation *r*^2^ are present, while after iterative filtering almost exclusively well-imputed SNPs are retained. On the contrary, filtering on the STITCH info score leads to inferior enrichment in well-imputed SNPs both at a 0.4 and at a 0.9 threshold. Note especially the large number of SNPs with very low ground truth correlation that are retained when filtering on the info score. The total number of SNPs is 6.42M for the original set, 5.39M for info score > 0.4, 3.2M for info score > 0.9, and 3.2M for the iterative filtering approach.

**Figure S5.**
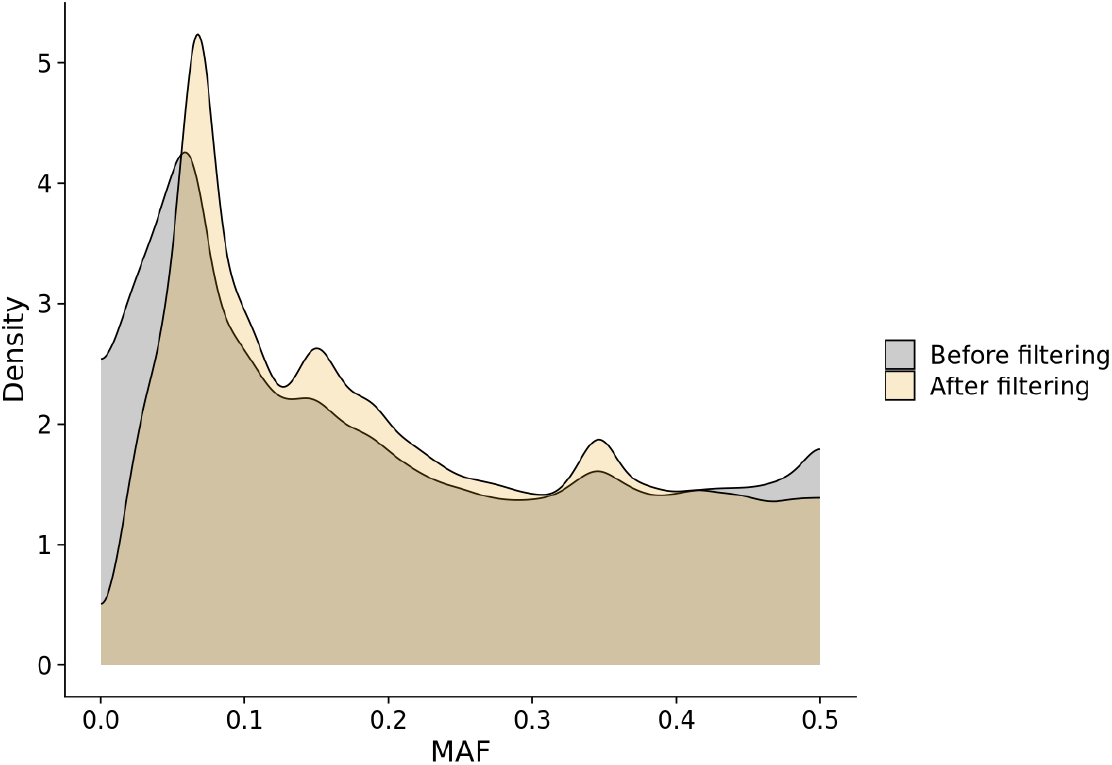
Distribution of the Minor Allele Frequency (MAF) of SNPs before and after the iterative filtering procedure described in this work. We observe that filtering preferentially retains SNPs with a MAF of approximately 0.06, while it depletes very rare or very common variants. A MAF of 0.06 is consistent with the frequency of a SNP being fixed in one founder line and absent from all the other founder lines when the line is used in 2 separate F2 crosses of 150 samples each (the most common scenario in our design). Non-variable SNPs (MAF = 0) were excluded.

**Figure S6.**
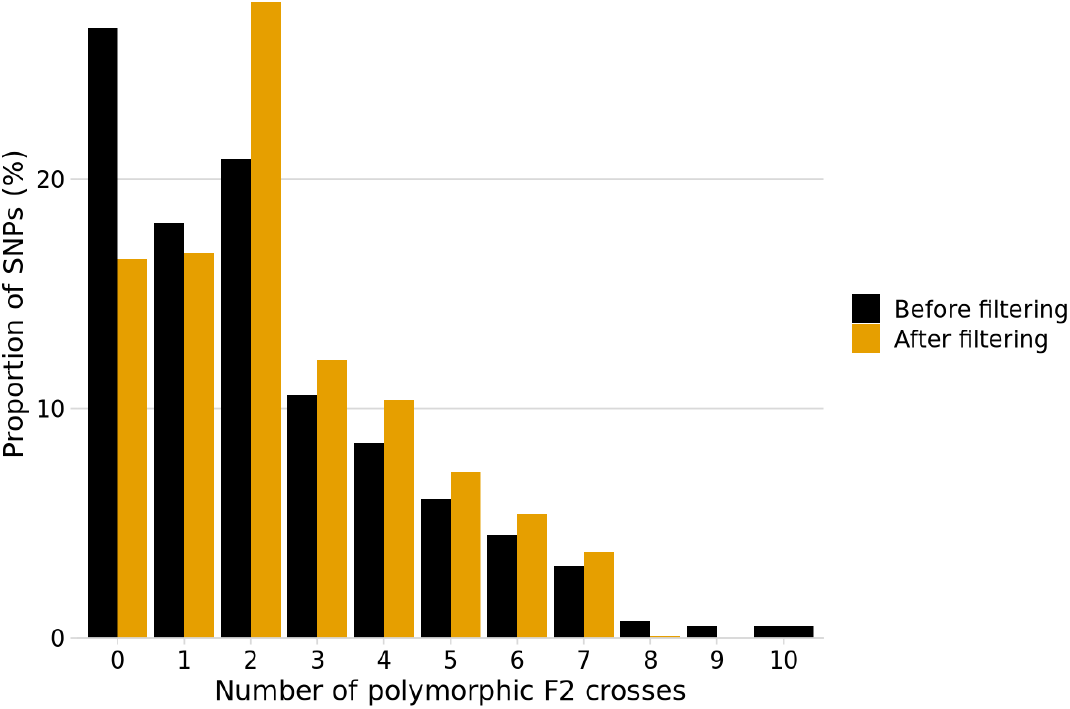
Strain distribution pattern for SNPs before and after the filtering procedure used in this work. We see an enrichment in SNPs segregating in 2 to 7 F2 crosses. Most founder lines are used in 2 separate F2 crosses in our design, while one founder line (72-2) is used in 7 crosses and one (22-1) is used in one cross only. We define a SNP as polymorphic in one cross when its minor allele frequency within the cross is greater than 0.4. Non-variable SNPs (MAF = 0 across all the samples) were excluded.

## References

Aida, T. (1921) ON THE INHERITANCE OF COLOR IN A FRESH-WATER FISH, APLOCHEILUS LATIPES TEMMICK AND SCHLEGEL, WITH SPECIAL REFERENCE TO SEX-LINKED INHERITANCE. Genetics, 6, 554–573.

Altshuler, D.M. et al. (2010) Integrating common and rare genetic variation in diverse human populations. Nature, 467, 52–58.

Bert, B. et al. (2016) Considerations for a European animal welfare standard to evaluate adverse phenotypes in teleost fish. EMBO J, 35, 1151–1154.

Bhattarai, G. et al. (2020) Genome Wide Association Studies in Multiple Spinach Breeding Populations Refine Downy Mildew Race 13 Resistance Genes. Frontiers in Plant Science, 11.

Blain, S.A. et al. (2024) Reduced hybrid survival in a migratory divide between songbirds. Ecology Letters, 27, e14420.

Broman, K.W. et al. (2019) R/qtl2: Software for Mapping Quantitative Trait Loci with High-Dimensional Data and Multiparent Populations. Genetics, 211, 495–502.

Browning, B.L. et al. (2018) A One-Penny Imputed Genome from Next-Generation Reference Panels. The American Journal of Human Genetics, 103, 338–348.

Cai, N. et al. (2015) Sparse whole-genome sequencing identifies two loci for major depressive disorder. Nature, 523, 588–591.

Danecek, P. et al. (2021) Twelve years of SAMtools and BCFtools. GigaScience, 10.

Davies, R.W. et al. (2021) Rapid genotype imputation from sequence with reference panels. Nat Genet, 53, 1104–1111.

Davies, R.W. et al. (2016) Rapid genotype imputation from sequence without reference panels. Nature Genetics, 48, 965–969.

Delaneau, O. et al. (2019) Accurate, scalable and integrative haplotype estimation. Nat Commun, 10, 5436.

Ewels, P.A. et al. (2020) The nf-core framework for community-curated bioinformatics pipelines. Nat Biotechnol, 38, 276–278.

Fitzgerald, T. et al. (2022) The Medaka Inbred Kiyosu-Karlsruhe (MIKK) panel. Genome Biology, 23.

Fuchsberger, C. et al. (2015) minimac2: faster genotype imputation. Bioinformatics, 31, 782–784.

Hanssen, F. et al. (2023) Scalable and efficient DNA sequencing analysis on different compute infrastructures aiding variant discovery. 2023.07.19.549462.

Hennig, B.P. et al. (2018) Large-Scale Low-Cost NGS Library Preparation Using a Robust Tn5 Purification and Tagmentation Protocol. G3 Genes|Genomes|Genetics, 8, 79–89.

Howie, B.N. et al. (2009) A Flexible and Accurate Genotype Imputation Method for the Next Generation of Genome-Wide Association Studies. PLOS Genetics, 5, e1000529.

Jaegle, B. et al. (2023) Extensive sequence duplication in Arabidopsis revealed by pseudo-heterozygosity. Genome Biology, 24, 44.

Kasahara, M. et al. (2007) The medaka draft genome and insights into vertebrate genome evolution. Nature, 447, 714–719.

Kluyver, T. et al. (2016) Jupyter Notebooks - a publishing format for reproducible computational workflows. In, Loizides, F.and Scmidt, B. (eds), Positioning and Power in Academic Publishing: Players, Agents and Agendas. IOS Press, Netherlands, pp. 87–90.

Lander, E.S. (1996) The New Genomics: Global Views of Biology. Science, 274, 536–539.

Li, N. and Stephens, M. (2003) Modeling Linkage Disequilibrium and Identifying Recombination Hotspots Using Single-Nucleotide Polymorphism Data. Genetics, 165, 2213–2233.

Li, Wenjie et al. (2024) Marker Density and Models to Improve the Accuracy of Genomic Selection for Growth and Slaughter Traits in Meat Rabbits. Genes, 15, 454.

Li, Yudong et al. (2022) Genetic parameters estimation and genome-wide association studies for internal organ traits in an F2 chicken population. Journal of Animal Breeding and Genetics, 139, 434–446.

Liu, S. et al. (2023) Utilizing Non-Invasive Prenatal Test Sequencing Data Resource for Human Genetic Investigation. 2023.12.11.570976.

McCarthy, S. et al. (2016) A reference panel of 64,976 haplotypes for genotype imputation. Nat Genet, 48, 1279–1283.

Mendel, G. (1866) Versuche über Plflanzenhybriden. Verhandlungen des naturforschenden Vereines in Brünn, Bd. IV für das Jahr 1865, 3–47.

Morgan, T.H. (1910) Sex Limited Inheritance in Drosophila. Science, 32, 120–122.

Nicod, J. et al. (2016) Genome-wide association of multiple complex traits in outbred mice by ultra-low-coverage sequencing. Nat Genet, 48, 912–918.

Pedersen, B.S. and Quinlan, A.R. (2017) Mosdepth: quick coverage calculation for genomes and exomes. Bioinformatics, 34, 867–868.

Picelli, S. et al. (2014) Full-length RNA-seq from single cells using Smart-seq2. Nat Protoc, 9, 171–181.

Poplin, R. et al. (2018) Scaling accurate genetic variant discovery to tens of thousands of samples. 201178.

R Core Team (2023) R: A Language and Environment for Statistical Computing R Foundation for Statistical Computing, Vienna, Austria.

Ribarska, T. et al. (2022) Optimization of enzymatic fragmentation is crucial to maximize genome coverage: a comparison of library preparation methods for Illumina sequencing. BMC Genomics, 23, 92.

Rubinacci, S. et al. (2021) Efficient phasing and imputation of low-coverage sequencing data using large reference panels. Nat Genet, 53, 120–126.

Scott, M.F. et al. (2021) Limited haplotype diversity underlies polygenic trait architecture across 70\hspace0.167emyears of wheat breeding. Genome Biology, 22.

The 1000 Genomes Project Consortium (2015) A global reference for human genetic variation. Nature, 526, 68–74.

Tommaso, P.D. et al. (2017) Nextflow enables reproducible computational workflows. Nature Biotechnology, 35, 316–319.

Van der Auwera, G.A. and O’Connor, B.D. (2020) Genomics in the Cloud O’Reilly Media, Inc.

Vasimuddin, Md. et al. (2019) Efficient Architecture-Aware Acceleration of BWA-MEM for Multicore Systems. In, 2019 IEEE International Parallel and Distributed Processing Symposium (IPDPS)., pp. 314–324.

Wang, D. et al. (2022) Cost-effectively dissecting the genetic architecture of complex wool traits in rabbits by low-coverage sequencing. Genetics Selection Evolution, 54, 75.

Wickham, H. (2016) ggplot2: Elegant Graphics for Data Analysis Springer-Verlag New York.

Wilke, C.O. (2020) cowplot: Streamlined Plot Theme and Plot Annotations for ‘ggplot2’.

Wittbrodt, J. et al. (2002) Medaka — a model organism from the far east. Nature Reviews Genetics, 3, 53–64.

Yao, E.J. et al. (2021) Systems genetic analysis of binge-like eating in a C57BL/6J x DBA/2J-F2 cross. Genes, Brain and Behavior, 20, e12751.

Zan, Y. et al. (2019) Genotyping by low-coverage whole-genome sequencing in intercross pedigrees from outbred founders: a cost-efficient approach. Genetics Selection Evolution, 51, 44.

Zha, C. et al. (2023) Combining genome-wide association study based on low-coverage whole genome sequencing and transcriptome analysis to reveal the key candidate genes affecting meat color in pigs. Animal Genetics, 54, 295–306.

